# Biosynthetic Gene Cluster Diversity Across Native and Introduced Populations of an Ectomycorrhizal Fungus, *Suillus luteus*

**DOI:** 10.1101/2024.09.16.612996

**Authors:** Brooke M. Allen, Cole Stevens, Rytas J. Vilgalys, Yi-Hong Ke, Milton Drott, Jason D. Hoeksema

## Abstract

Non-pathogenic fungi are being increasingly recognized for their involvement in biological invasions, where they have the potential to cause ecological and economical harm. One notable example is the ectomycorrhizal (EcM) fungus, *Suillus luteus*, which is commonly found co-invading with non-native pines that have escaped from commercial pine plantations across the Southern Hemisphere. In such invasions, selection pressures imposed by novel plants, soil microbes, and environmental conditions, as well as impoverished assemblages of co-invading EcM fungi, may exert particularly strong selective effects and contribute to driving evolutionary divergence in traits. We investigated the potential impact of this global co-introduction on biosynthetic gene cluster (BGC) diversity in native and introduced populations of *S. luteus*. We found that native populations demonstrated higher BGC diversity at both the gene and clan levels compared to introduced populations. Additionally, we identified a collection of unique BGCs as well as 24 highly conserved clans, including two featuring known pathways, within both native and introduced populations. Furthermore, our study confirmed the presence of three previously identified clades of *S. luteus* with distinct gene cluster family (GCF) diversity. Thus, our study concludes that the introduction of *S. luteus* has driven evolutionary changes in its BGCs, underscoring the broader implications of global co-introductions on fungal adaptation in new environments.

## 1. Introduction

Mycorrhizae are classic examples of microbial symbiosis, wherein fungi establish associations with plant roots, facilitating the exchange of soil nutrients (such as nitrogen and phosphorus) for photosynthetically fixed carbon (Smith and Read, 2010). Ectomycorrhizal (EcM) fungi are commonly found in association with dominant trees and are integral members of rhizosphere communities in temperate and boreal forests. They play essential roles in facilitating seedling establishment, promoting tree growth, and influencing forest succession dynamics (Tedersoo et al., 2010; Kałucka and Jagodziński, 2017). EcM fungi are also being increasingly recognized for their involvement in biological invasions, often due to their tendency to co-invade with invasive plant species into native ecosystems, where they have the potential to cause ecological and economical harm (Vellinga et al., 2009; Dickie et al., 2010; Hayward et al., 2015).

One notable example is the fungal genus *Suillus* (phylum *Basidiomycota*: subphylum *Agaricomycotina*: Class *Boletales*), which is ubiquitously found co-introduced and co-invading with non-native species of *Pinaceae* and is an emerging model system for studying the ecology and evolution of EcM interactions (Lofgren et al., 2024). *Suillus luteus* is one of the most widely introduced species of EcM fungi (Policelli et al., 2019), and has been found to be a major facilitator of pine invasions outside of plantations (Hayward et al., 2015). Native to Eurasia, *S. luteus* was introduced to the Southern Hemisphere and North America approximately 130-150 years ago (Peck, 1887; Vellinga et al., 2009) alongside obligately mycorrhizal pine trees (Burdon and Chilvers, 1977; Richardson et al., 1994). Across its introduced range, *S. luteus* associates with many different novel pine hosts such as *Pinus radiata, P. resinosa, P. strobus, P. contorta, P. patula*, and *P. ponderosa* (Vellinga et al., 2009; Nguyen, 2016; Policelli et al., 2019). A phylogenetic analysis by Ke et al. (2024, unpublished manuscript) of *S. luteus* SNPs extracted from the same genomic sequences we analyze here, identified three distinct, deeply divergent clades: Asia, Northern Europe, and Central Europe. Each clade exhibited unique geographic distribution patterns, with clade Asia forming a sister group to clade Northern Europe. They found that all individuals from introduced populations, except North America, had detectable differentiation from the Central Europe source population, indicating that evolution is occurring in introduced populations.

The repeated global co-introduction of *S. luteus* and its pine hosts provides an opportunity for exploring rapid evolution, co-evolution, and local adaptation of mycorrhizal traits in both plants and fungi (Hoeksema et al., 2020). In this study we focus specifically on biosynthetic gene clusters (BGCs), which are clustered groups of genes involved in natural product (also known as secondary metabolite) pathways in the genomes of bacteria, plants, and fungi. Unlike primary metabolites, natural products are considered non-essential for growth and development. Instead, they are thought to provide selective advantages by performing ecological functions that are still widely unknown, although many may play fundamental roles in diverse species interactions as chemical weapons and signals (Walsh and Tang, 2017; Keller, 2019; Mudbhari et al., 2023; Zhang et al., 2024). Just as ecologically relevant animal traits like teeth or claws evolve, so do the biochemical traits of fungi. The genetic diversity of BGCs, often linked to structural differences in molecular products (Medema et al., 2014), can result in variation in biological activities and subsequent changes in ecological interactions. In addition, there is a high cost associated with maintaining BGCs, and thus the strength of selection on them is likely also high (Fischbach et al., 2008). However, despite the potential for introduced microbial populations to evolve rapidly in response to new environments (Osbourn, 2010; Colautti and Lau, 2016), we still lack understanding of how these rapid changes affect the molecular traits of introduced microbes, including EcM fungi.

Therefore, in this study we aimed to a) survey BGC diversity across native and introduced populations of *S. luteus*, b) determine if there are biogeographical patterns in BGC distribution among populations, and c) evaluate evolutionary relationships among BGCs of *S. luteus*. To address these objectives, we identified putative BGCs in 258 *S. luteus* genomes from native and introduced populations, constructed BGC similarity networks, and grouped similar BGCs into gene cluster families (GCFs) and then similar GCFs into clans.

## 2. Materials and Methods

### 2.1. Dataset

258 *Suillus luteus* whole genome Illumina sequences (15.3x to 67.1x) (Table 1, supplementary table 2, supplementary figure 1) were downloaded from the Genome Portal of the Department of Energy Joint Genome Institute (Nordberg et al., 2014; Nguyen, 2016) and preprocessed using Trimmomatic (v0.39) (Bolger et al., 2014) to remove adapter sequences, trim low-quality leading and trailing bases (threshold: 20), apply sliding window quality trimming (window size: 4 bases, mean quality threshold: 25), and discard reads shorter than 50 bases after trimming. FastQC (v0.12.1) was used to verify the quality of the raw sequences both before and after trimming. Denovo genome assembly was then performed with SPADES (v3.11.1) (Bankevich et al., 2012) followed by gene annotation using Augustus (v3.5.0) (Stanke and Morgenstern, 2005). The quality of the assemblies was verified using BUSCO (v5.6.1) (Manni et al., 2021) to assess BUSCO scores and n50 (supplementary table 1). System details for the work conducted in this study are as follows: Python version 2.7.18 and OS Linux-4.12.14-122.150-default-x86_64-with-SuSE-12-x86_64. For additional information and to access the code utilized in this study, please refer to the provided [GitHub repository](https://github.com/ballen3/All_phd_ch_1_code).

**Table 1.** Country of origin, sample size, and status of the 258 *Suillus luteus* whole genome Illumina sequences.

| Country | Sample Size | Status |
| --- | --- | --- |
| Austria | 8 | Native |
| Belgium | 62 | Native |
| Estonia | 11 | Native |
| France | 1 | Native |
| Germany | 1 | Native |
| Italy | 2 | Native |
| Japan | 14 | Native |
| Norway | 10 | Native |
| Russia | 2 | Native |
| Sweden | 1 | Native |
| Taiwan | 1 | Native |
| Argentina | 20 | Introduced |
| Australia | 32 | Introduced |
| Canada | 4 | Introduced |
| Chile | 17 | Introduced |
| New Zealand | 31 | Introduced |
| Tanzania | 1 | Introduced |
| USA | 40 | Introduced |

### 2.2. Data analysis

BGCs were predicted from the 258 assembled *Suillus luteus* genomes using AntiSMASH (v7.1.0) (Medema et al., 2011) with the fungal taxon flag and all extra features on (cb-general, cb-knownclusters, cb-subclusters, pfam2go, and clusterhmmer) (supplementary table 2). Similarity networks and gene cluster families (GCFs) were then generated using BiG-SCAPE (v1.1.5) (Navarro-Muñoz et al., 2020) with the no hybrids, mix, mibig, and singleton parameters selected and the default cutoff of 0.3 similarity score. Resulting similarity networks were rendered with Cytoscape and analyzed with Cytoscape’s Network Analyzer (v3.10.1) as undirected graphs (Shannon et al., 2003). RStudio (2023.12.0+369 “Ocean Storm”) was used for all statistical analyses and plot building (R Packages: ggplot2, ggcorrplot, dplyr, tidyverse, vegan, and ellipse). GCFs are groups of BGCs that have similar protein domain content, order, copy number, and sequence identity. Clans (groups of similar families) were identified from the BiG-SCAPE network visualization of clustered GCFs (as seen in Figure 1).

**Figure 1.**
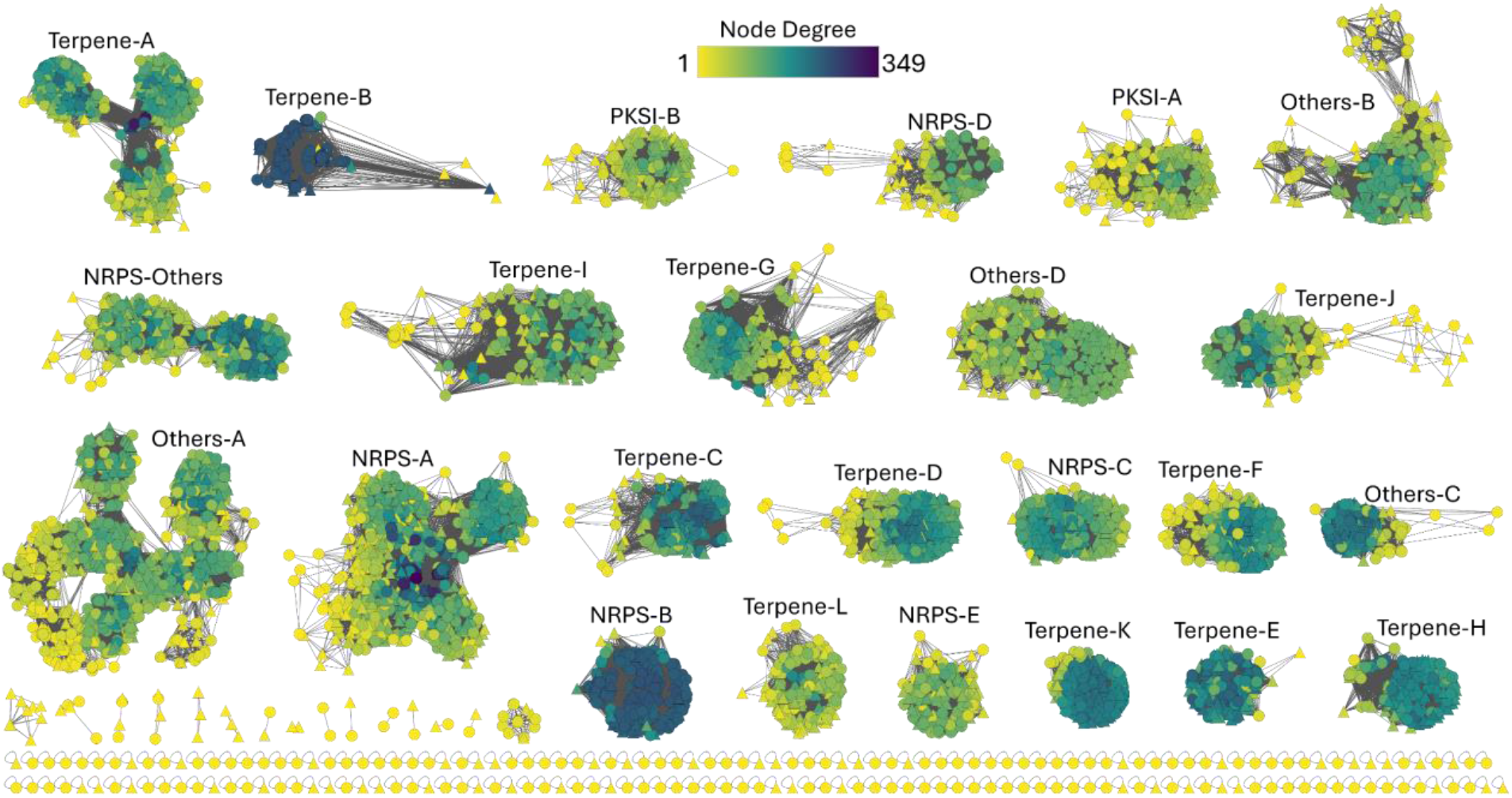
Network visualization of clustered BGCs. Nodes represent individual BGCs and edges represent similarity between nodes. Node positions reflect similarities, with closer nodes indicating higher similarity based on dynamic similarity scaling (DSS). Edge lengths are proportional to the raw DSS non-anchor values. Color gradient depicts node degree (Yellow=low, Blue=high degree) and node shape represents status (circle=Introduced, triangle=Native).

An approximate Wilcoxon rank sum test was used to identify clans where status (native and introduced) had a significant effect on degree, which is the number of connections made by nodes in the BiG-SCAPE network. From each of these clans, three levels of degree (high, medium, and low) were independently determined by creating three roughly even groups across the range of degrees in each clan. From each of those degree levels, five BGCs were randomly sampled for each selected clan and visualized with Clinker (v1.0) (Gilchrist and Chooi, 2021) via the CompArative GEne Cluster Analysis Toolbox (CAGECAT) web server. To compare the average number of BGCs per genome between groups, we applied the Wilcoxon rank-sum test/Mann-Whitney U-test with continuity correction. A 2-proportion Z-test was used to compare the proportions of BGC classes between native and introduced populations. Principal Coordinate Analysis (PCoA) was conducted to investigate the similarity of GCF composition (presence/absence with Jaccard Index) among genomes across three clades identified by Ke, et al. (2024, unpublished manuscript), Asia (Japan and Taiwan), Northern Europe (Austria, Norway, and Russia), and Central Europe (All other Countries). Bootstrapping (R=10,000) was employed to estimate the proportions of native, introduced, and shared GCFs, as well as the proportion of singletons. Additionally, a combination of Welch’s t-test and Mann-Whitney U-test was conducted to determine the effect of status on gene group count/BGC for each clan, with data transformations performed as necessary.

## 3. Results

### 3.1. BGC Diversity and Distribution

AntiSMASH predicted 8,238 putative BGCs across the 258 genomes, including three BGCs with sequence similarity (0.3) from the Minimum Information about a Biosynthetic Gene cluster (MiBIG) repository. BiG-SCAPE sequence similarity network analysis of the predicted BGCs resulted in the identification of 277 distinct gene cluster families (GCFs) and 181 singletons (unique nonclustered BGCs). Further similarity-based clustering of the 277 non-singleton GCFs resulted in 39 clans, each representing groups of similar GCFs (Figure 1). Of these 39 clans, 24 included more than 100 BGCs and are hereafter referred to as ‘large’ clans. Each large clan was assigned a name based on its BGC class, followed by a letter for differentiation (eg. Terpene-A, Terpene-B, ect.).

There was no significant difference observed in the number of BGCs per genome between introduced (North and South America, Oceania, and Africa; see Table 1 for Countries) (average of 32.19 per genome) and native (Europe and Asia; see Table 1 for Countries) (average of 31.60 per genome) populations (U = 8554, p = 0.54). However, there were significant differences in the proportion of GCFs for some classes of BGCs between native and introduced populations, with the introduced populations exhibiting a significantly higher proportion of ribosomally synthesized and post-translationally modified peptides (RiPPs) (X^2^ = 5.554, df = 1, p= 0.018), while the native populations had a significantly higher proportion of terpenes (X^2^ = 4.4833, df = 1, p = 0.034) (Figure 2a). NRPS were the most prevalent class of BGC by number of GCFs predicted across all genomes, comprising 31% of all GCFs. This was followed by “Others” (28%) and Terpenes (28%), then RiPPs (9%), PKSI (4%), hybrid PKS-NRPs (0.4%) and PKSother (0.4%) (Figure 2b). The majority of BGCs classified as “Other” consisted of fungal-ripp-like product predictions. In addition, hybrid PKS-NRPs and “PKSother” were exclusively found in introduced populations.

**Figure 2.**
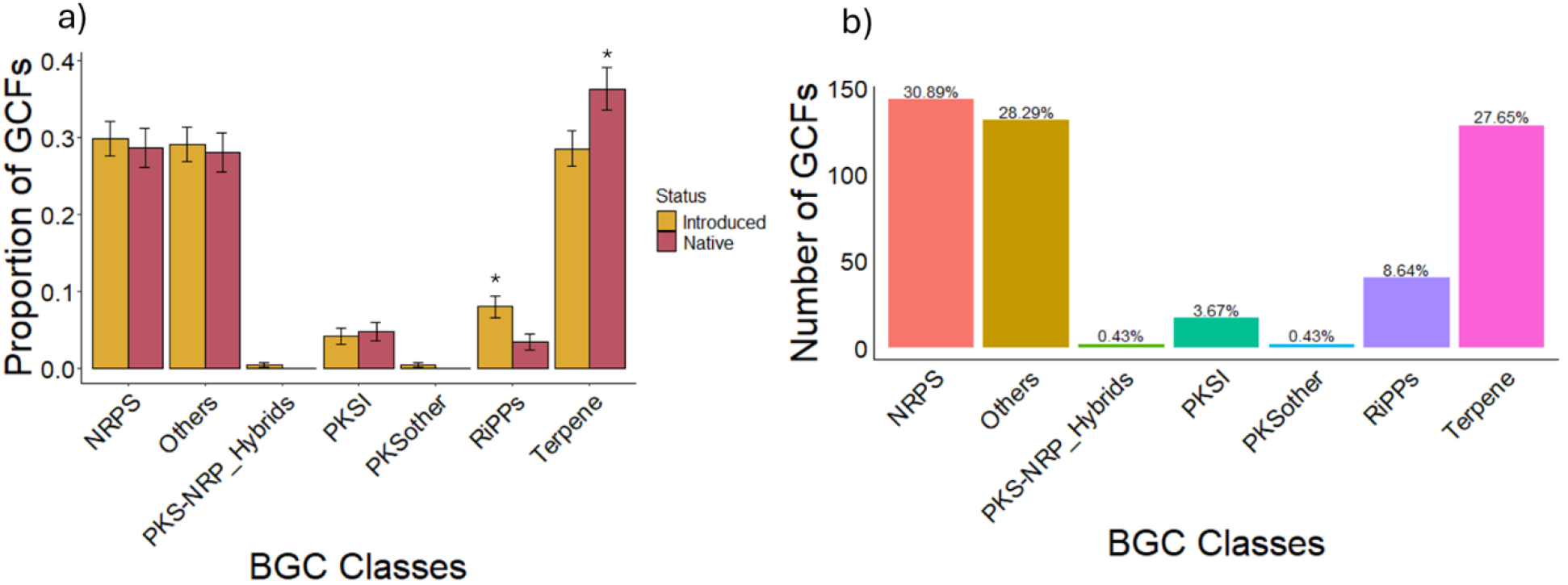
Visualizations of GCFs across all BGC classes. (a) Proportion of GCFs compared between native and introduced populations, with error bars representing standard error and asterisks indicating a significance level of ≤0.05. (b) Total abundance (counts) of GCFs from each BGC class, with percentages above each bar representing the overall percent abundance of each BGC class, without distinction between native and introduced populations.

### 3.2. GCF Diversity and Biogeographical Distribution

None of the identified GCFs contained BGCs from all 258 genomes. However, we observed that in six of the GCFs (GCF 3071 (87.7%), GCF 5626 (69.76%), GCF 6141 (56.20%), GCF 492 (55.81%), GCF 2040 (51.93%), and GCF 7163 (50.77%)), at least 50% of the sampled genomes contributed one or more BGCs. The GCF with the greatest genomic representation was a terpene, which contained BGCs from 87.7% of all genomes. In addition, four clans (terpeneA, terpeneB, nrpsA, and othersA) did contain BGCs from all 258 genomes, or 100% BGC representation. Three more clans (nrps_others, terpeneC, othersC) contained BGCs from >99% of genomes and 8 more (othersD, nrpsB, terpeneE, othersB, terpeneD, terpeneF, nrpsC, terpeneG, and terpeneH) contained BGCs from 80-98% of genomes. None of the 24 large clans contained less than 61% of BGCs from all genomes.

The distribution of GCFs across populations was examined using bootstrapping (R=10000). Approximately 16% (95% CI: 12.11% - 20.85%) of non-singleton GCFs within the large clans were uniquely associated with native populations, paralleled by an equivalent proportion of 16% (95% CI: 12.29% - 20.83%) exclusively linked to introduced populations (Figure 3a). Approximately 67% (95% CI: 62.24% - 71.92%) of GCFs were found to be shared between both native and introduced populations (Figure 3a). Singletons comprised 5.1% (95% CI: 3.2%-6.9%) of identified GCFs in native populations and 14.0% (95% CI: 11.1%-16.9%) in introduced populations (Figure 3b). The overall proportion of singletons across the entire dataset was 25.4% (95% CI: 22.3%-28.6%).

**Figure 3.**
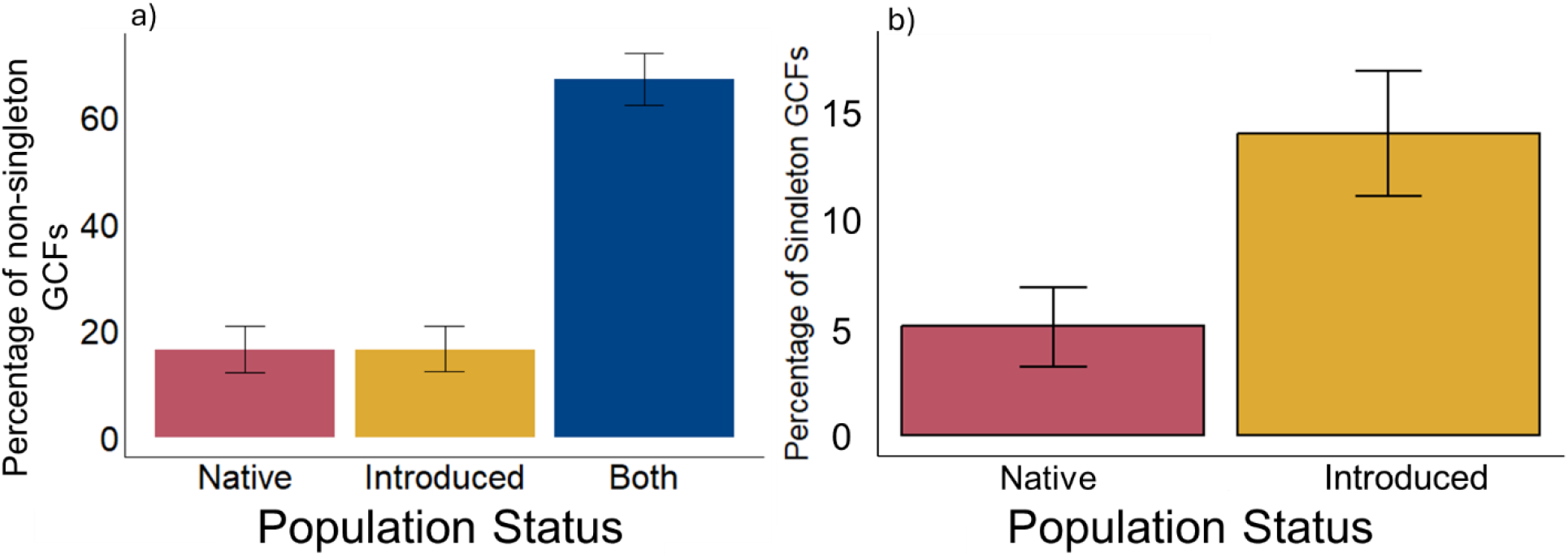
Percentage of a) non-singleton GCFs uniquely associated with native and introduced population and those shared between both types of populations and b) percentage of singleton GCFs associated with native and introduced population. Error bars represent bootstrapping CIs (95% CI).

PCoA was performed using the Jaccard Index calculated from the presence-absence data of GCFs across all genomes to visualize compositional differences. The first two principal coordinates explained 4.89% and 2.99% of the total variance, respectively. The cumulative variance explained by the first three axes was 10.51%. The PCoA plot shows that the Asia clade is mostly distinct but overlaps with Northern Europe, while Northern Europe itself overlaps with both Asia and Central Europe. Central Europe is made up of both native and introduced statuses, with introduced forming a tightly clustered group nested within the more dispersed native group (Figure 4). Central European points clustered with Asia and Northern Europe originated from Germany and Italy (native), as well as from Argentina and Chile (introduced).

**Figure 4.**
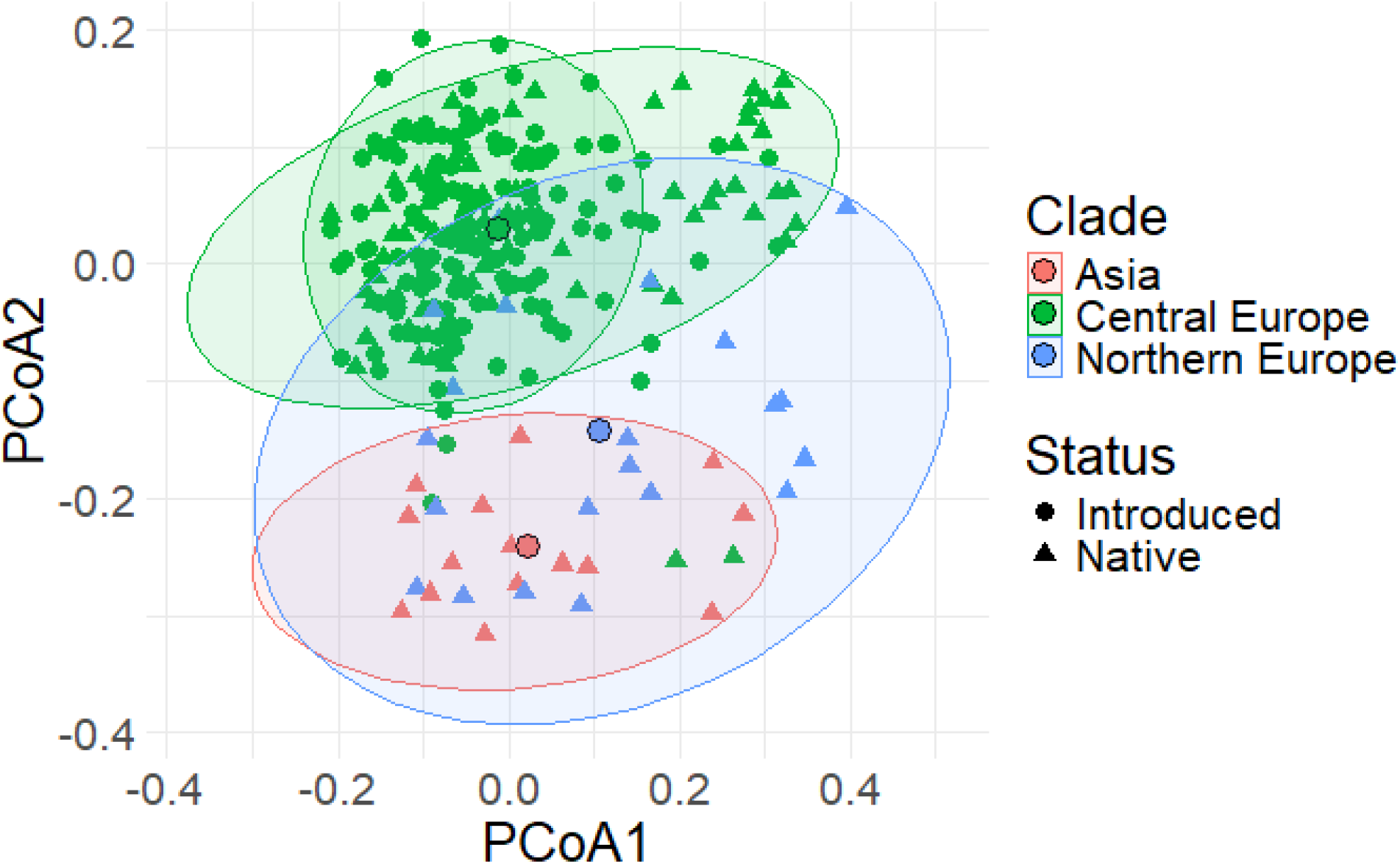
PCoA plot of overall GCF composition within genomes (presence/absence), colored by clade as defined by Ke et al. (2024, unpublished manuscript). Ellipses represent 95% confidence intervals for both clade and status within clade groupings where both statuses are present.

Three annotated BGCs sourced from the MiBiG repository were identified. These BGCs demonstrated sequence similarity with several BGCs predicted from *Suillus luteus* genomes. Specifically, two NRPS-like BGCs, namely atromentin BGC (BGC0002277) from *Suillus grevillei* and enterobactin BGC (BGC0002476) from *Escherichia coli* str. K-12 substr. MG1655, along with one terpene, (+)-δ-cadinol BGC (BGC0002708) from *Coniophora puteana* RWD-64-598 SS2, were identified. The atromentin and (+)-δ-cadinol BGCs were found within GCFs that encompassed a substantial number of BGCs, totaling 953 and 255, respectively. In contrast, the enterobactin BGC was part of a smaller GCF, containing only one additional BGC.

### 3.3. BGC Diversity within Clans and Evolution within BGCs

We assessed BGC diversity and evolution by examining variation within clans, which was determined based on degree of connectivity and the number of gene groups per BGC. In six clans, status (native and introduced) significantly influenced the degree of connectivity. In five of those clans, degree was higher in native than in introduced BGCs: PKS1A (U=3736.5, p=0.010), TerpeneD (U=6234.5, p=0.0027302731), TerpeneF (U=5693.0, p=0.00037), TerpeneH (U=5646.0, p=0.031), and TerpeneK (U=4523.0, p=0.0028). TerpeneE had higher degree in introduced BGCs compared to native (U=9969.0, p=0.00047).

From each of those six clans, we randomly selected 15 BGCs (5 BGCs per degree level of high, medium, and low) to compare via Clinker visualization. As expected, we observed gene groups that were shared across all BGCs within a clan. Additionally, we found that BGCs within each clan showed notable differences in gene size, gene copy number, gene group diversity, and sequence similarity (Figure 5).

**Figure 5.**
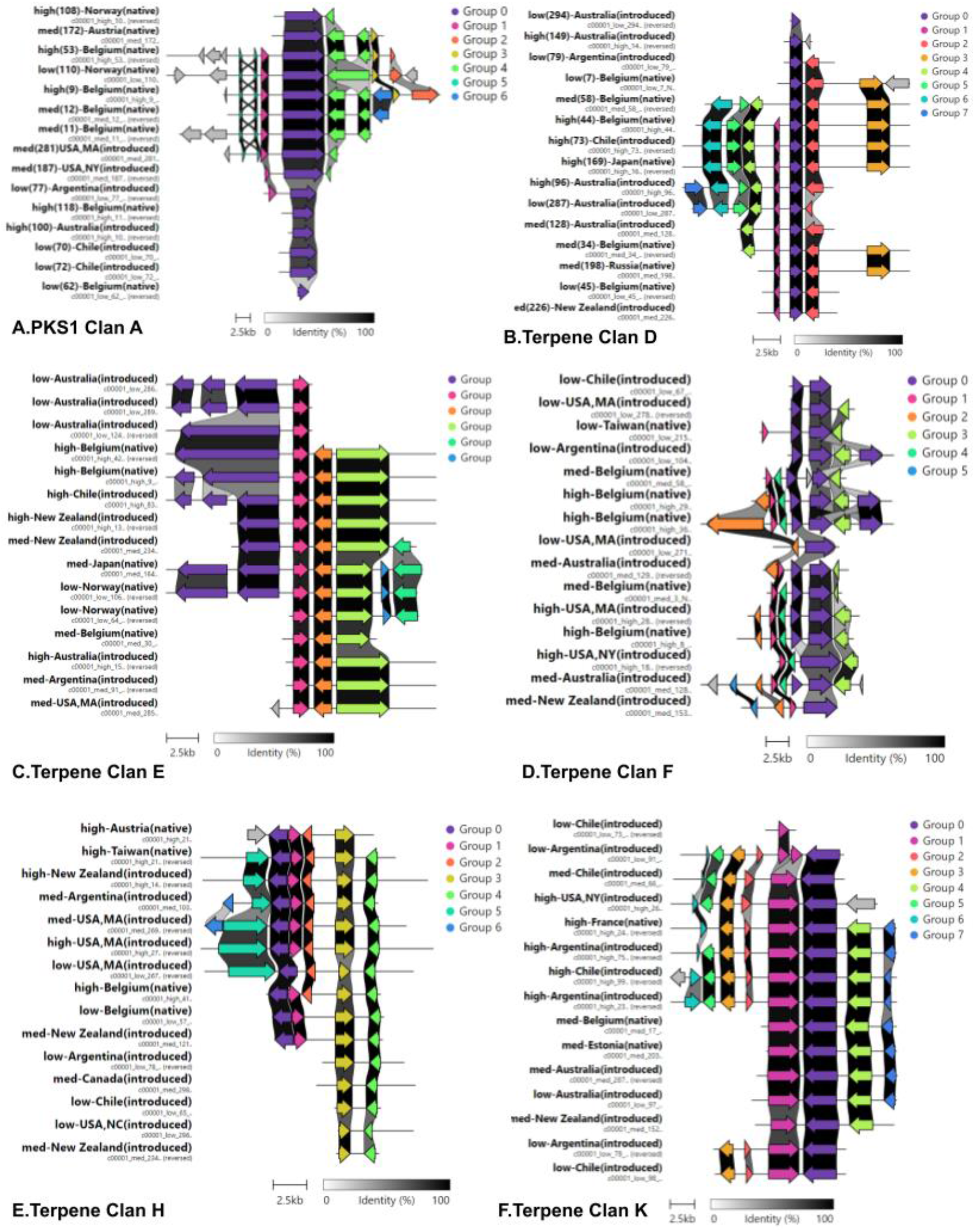
Clinker visualization of BGCs from three different degree levels within the six clans having a significant effect of status on degree.

To further understand BGC diversity within clans, we analyzed the number of gene groups per BGC between native and introduced populations in all 24 large clans. There was a significant difference in gene group count per BGC between the native and introduced populations across 12 clans (refer to Table 2 for statistical values), with native BGCs exhibiting a higher gene group count per BGC compared to introduced BGCs in all instances (Figure 6).

**Table 2.**
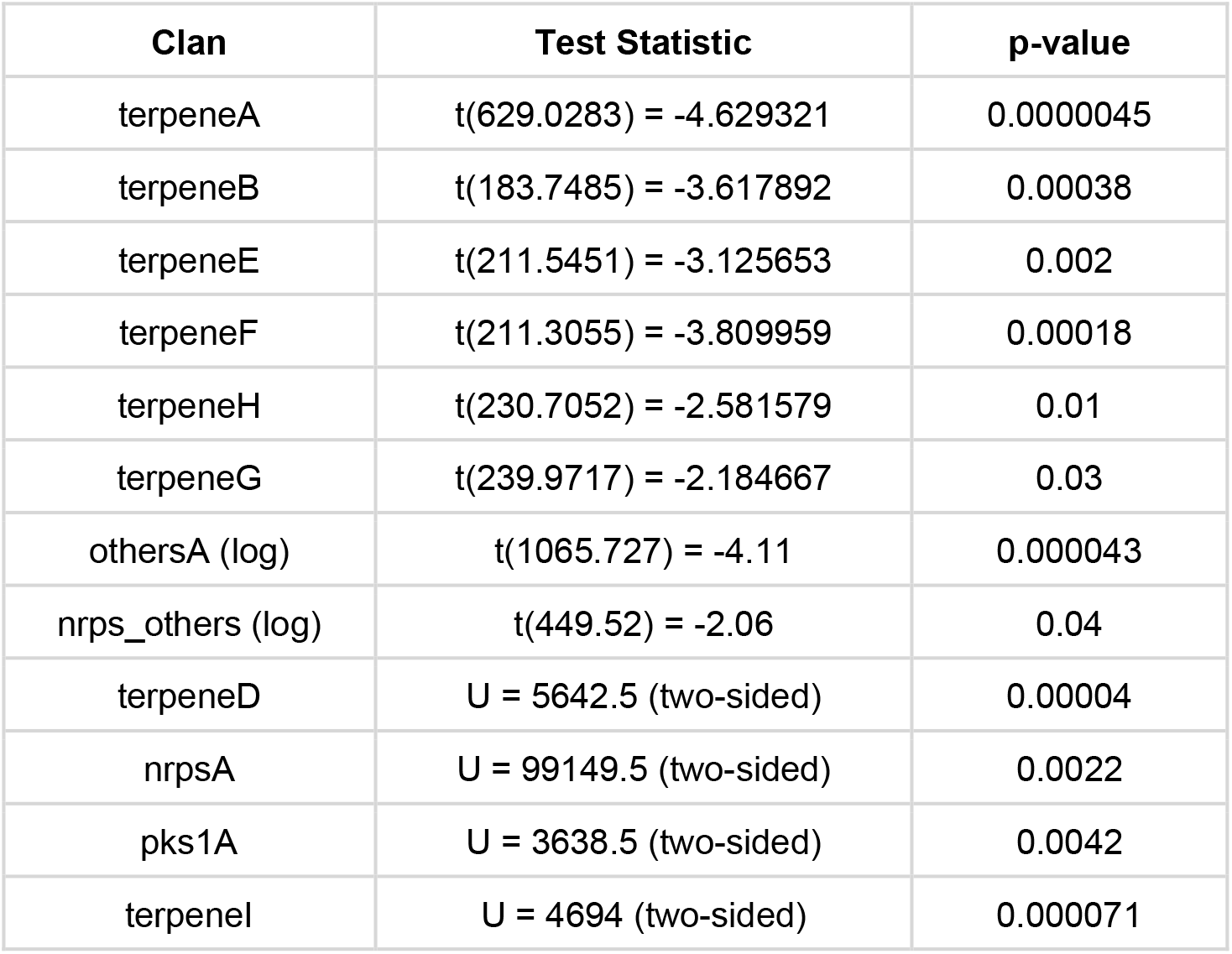
Statistical values for clans with significant differences between native and introduced populations for gene group count per BGC.

| Clan | Test Statistic | p-value |
| --- | --- | --- |
| terpeneA | t(629.0283) = -4.629321 | 0.0000045 |
| terpeneB | t(183.7485) = -3.617892 | 0.00038 |
| terpeneE | t(211.5451) = -3.125653 | 0.002 |
| terpeneF | t(211.3055) = -3.809959 | 0.00018 |
| terpeneH | t(230.7052) = -2.581579 | 0.01 |
| terpeneG | t(239.9717) = -2.184667 | 0.03 |
| othersA (log) | t(1065.727) = -4.11 | 0.000043 |
| nrps_others (log) | t(449.52) = -2.06 | 0.04 |
| terpeneD | U = 5642.5 (two-sided) | 0.00004 |
| nrpsA | U = 99149.5 (two-sided) | 0.0022 |
| pks1A | U = 3638.5 (two-sided) | 0.0042 |
| terpenel | U = 4694 (two-sided) | 0.000071 |

**Figure 6.**
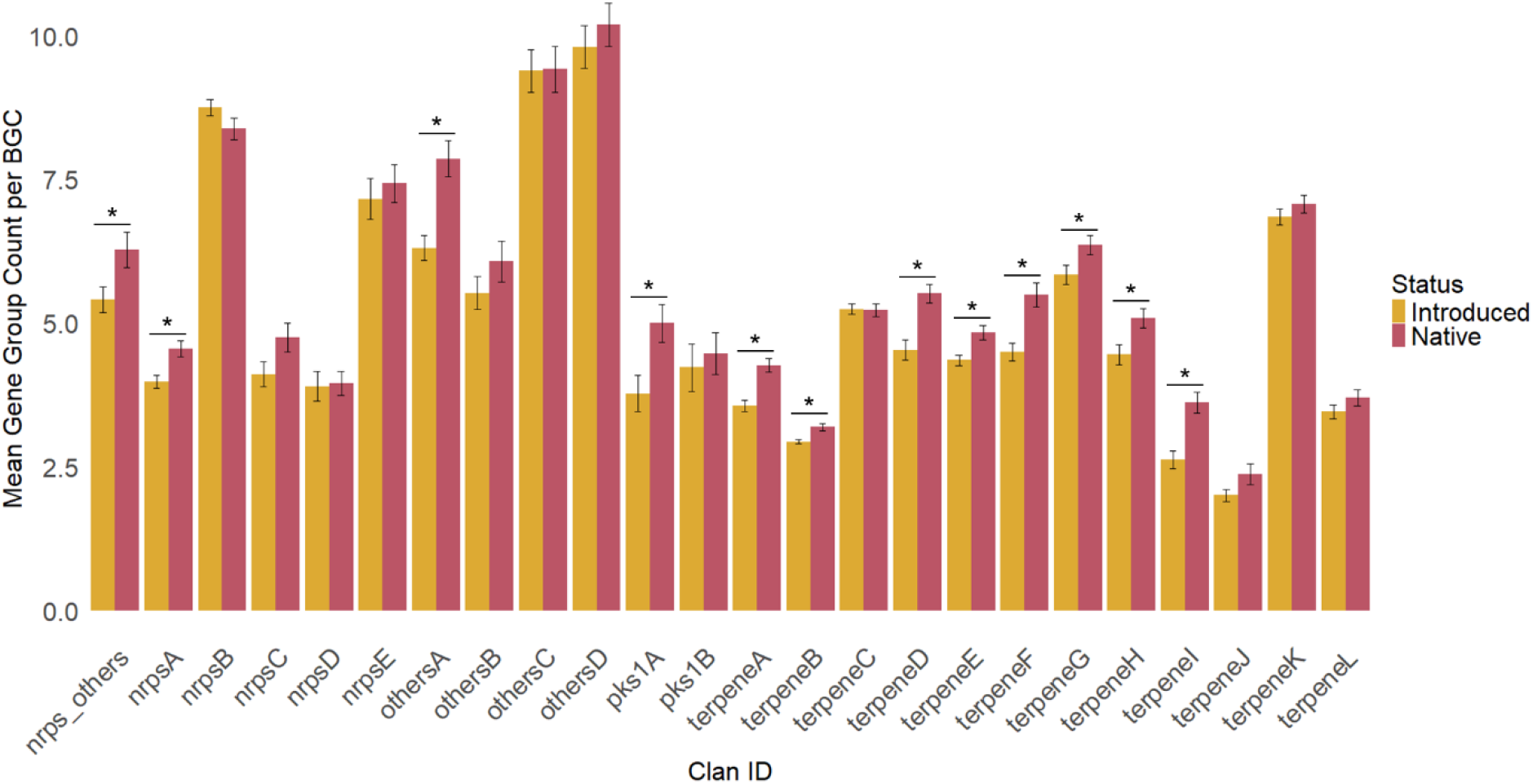
Mean gene group count per BGC for each clan with more than 100 BGCs is depicted, with error bars representing standard error. Asterisks denote significant differences (p<0.05).

## 4. Discussion

This study aimed to explore BGC diversity and evolution in an ectomycorrhizal fungus by surveying BGC diversity across native and introduced populations of *S. luteus*, examining biogeographical patterns in BGC distribution, and evaluating evolutionary relationships among *S. luteus* BGCs. Employing a hierarchical clustering approach, we organized similar BGCs into GCFs and further grouped these into clans. This method enabled us to explore biosynthetic diversity across various scales, from individual genes to groups of populations, offering a comprehensive perspective on this diversity.

We found that there was high overall biosynthetic potential, with both native and introduced populations producing approximately 32 BGCs per genome on average, which is similar, although slightly lower, to what has been found in *S. hirtellus* and *S. decipiens* (Mudbhari et al., 2023). As found previously in other species of *Suillus* (Mudbhari et al., 2023; Lofgren et al., 2024), terpenes and NRPSs were, proportionally, highly prevalent classes among GCFs overall, but unlike previous studies, we found that the class identified as “Others” by AntiSMASH was also among the most prevalent classes of GCFs (Figure 2b). Notably, the Other classification consisted mainly of fungal-ripp-like product predictions. Terpenes are the largest class of putative natural products and are widespread in both plants and fungi (Gershenzon and Dudareva, 2007; Plett et al., 2024), and while in many cases the ecological roles of microbially produced terpenes are unknown, many have been found to play important roles in competition and communication (recognition and response) with hosts and other microbes (Lofgren et al., 2021; Wu et al., 2022; Mudbhari et al., 2023). RiPPs represent another significant class of fungal natural products and, like terpenes, have been associated with competitive interactions and communication (Li and Rebuffat, 2020;

Ford et al., 2022). RiPPs are also responsible for producing toxins such as amatoxins and phallotoxins from *Amanita phalloides*, which are often associated with defense against consumption, but may also play a role in the control of fungal cell growth at reduced quantities (Ford et al., 2022). As for NRPSs, their most well-known ecological role is in relation to siderophores, which are involved in plant and animal virulence and iron homeostasis (Bills et al., 2014).

Proportionally, native populations had more terpene GCFs compared to introduced populations, while introduced populations exhibited more RiPPs compared to the native populations (Figure 2a). Given the consistent proportions of other classes across populations, these differences are likely due to selective pressures related to differences in environmental factors between the introduced and native ranges, rather than being a result of genetic drift alone. In the introduced range, the EcM fungal community exhibits considerably lower diversity compared to native ranges (Policelli et al., 2019; Hoeksema et al., 2020), potentially leading to different competitive pressures that make the production of some terpenes disadvantageous and some RiPPs advantageous.

Additionally, we observed a high level of conservation among large clans, with all 24 large clans containing BGC representation from >60% of all genomes. Clans encompass multiple GCFs, grouping together BGCs that, while structurally and functionally related, can still exhibit significant variation. Four clans, including two terpenes, one NRPS, and one Other, were conserved across all 258 genomes. We also see this conservation reflected in the high proportion of GCFs shared across both native and introduced populations (67% of non-singleton GCFs) (Figure 3a). Together, these results indicate that at the broad level of functional diversity there is a core set of conserved BGCs, especially terpenes, NRPS, and Others/fungal-ripp-like product predictions. In contrast, we also observed that unique, unclustered (singleton) BGCs are common, constituting approximately a quarter of all GCFs, with the introduced populations exhibiting a higher proportion of these singleton BGCs compared to the native populations (Figure 3b). These BGCs are unlike any others predicted in our analysis or found in the MiBIG database, suggesting that local adaptation, particularly in the case of introduced populations, may be generating functionally novel BGCs.

Two highly conserved clans contained known annotated BGCs, atromentin and (+)-δ-cadinol respectively, sourced from *Suillus grevillei* and *Coniophora puteana* in the MiBIG repository (Terlouw et al., 2023). The identification of known BGCs within these clans provides us with the benefit of understanding the general chemical structure for BGCs within those clans and allows us to speculate on the role of those BGCs in an ecological context based on previous work. Atromentin, an NRPS BGC and terphenylquinone natural product, is chemo-taxonomically connected to Boletales and produces a precursor pigment that forms the basis of hundreds of other pigments, which give the fruiting bodies of boletes their color (Braesel et al., 2015; Tauber and Hintze, 2020; Seibold et al., 2023). Atromentin and its derivatives have been linked to inhibition of bacterial motility, modulation of biofilms, and reduction of Fe3+ during Fenton based decomposition of organic material (Braesel et al., 2015; Seibold et al., 2023). In addition, the atromentin biosynthetic pathway independently evolved twice in basidiomycetes, which implies significant ecological importance (Seibold et al., 2023). (+)-δ-cadinol (also known as “torreyol” or “pilgerol) is a cadinane-type sesquiterpene, which as a family is generally associated with decay resistance and antifungal activity (Wu et al., 2005; Ringel et al., 2022). (+)-δ-cadinol has been extensively studied from a pharmaceutical and biotechnology perspective, but there is a lack of studies that focus on determining its ecological role.

Biogeographically, there are three distinct groups based on GCF composition (presence/absence) within genomes (Figure 4). These groups align with the three deeply diverged clades identified by Ke et al. (2024, unpublished manuscript), and therefore support the existence of 3 distinct populations of *S. luteus*. The separation of clades Asia and Northern Europe indicate they have distinct GCF (and therefore also BGC) compositions compared to clade Central Europe. Curiously, the Northern Europe clade contains genomes from individuals that are not geographically separated from those in Central Europe, while also being highly geographically separated from those in Asia. This observation raises the possibility of a cryptic *Suillus* species in Asia and Northern Europe as an explanation, rather than reproductive isolation among populations of *S. luteus* (Ke et al., 2024, unpublished manuscript). We also found that all introduced genomes were nested within the Central Europe clade group, with much less dispersion compared to the native Central Europe genomes. This observation supports the hypothesis that all introduced populations originated from Central Europe, and also indicates that introduced populations are less differentiated in their BGC diversity from each other than from the Northern Europe and Asia clades.

While the average degree of connectivity was not different between native and introduced populations overall, five clans exhibited significantly higher connectivity of native BGCs. This result suggests that native BGCs within these clans are more interconnected, indicating higher redundancy or greater diversity among closely related BGCs. This increased connectivity may be due to a longer evolutionary history and more gene flow among native populations, allowing for more related BGC variants to develop. In contrast, introduced BGCs within these clans show fewer variants or have undergone adaptations that resulted in less similar, less connected BGCs. Novel BGCs often emerge from existing ones, although *de novo* formation is also possible (Rokas et al., 2020). One introduced clan had a significantly higher degree of connectivity among introduced BGCs, which may indicate rapid diversification or retention of diversity from the original native population.

Visualizations of BGCs, randomly selected by degree level from clans with significantly different degrees, revealed evolutionary diversification within clans and the genetic connections between them. BGCs within the same clan shared common gene groups but varied in their architecture and sequence similarity (Figure 5). Notably, 12 clans had a significantly higher number of gene groups per BGC in native populations compared to introduced populations, indicating a more complex and diverse biosynthetic landscape with more genetic exchange (Figure 6). This, along with our degree of connectivity trends, suggests that introduced populations have less divergent BGCs within these clans compared to native populations. Lower BGC divergence in introduced habitats may result from selective pressures favoring the loss or reduction of certain gene groups deemed non-essential or disadvantageous in those novel habitats (Fischbach et al., 2008; Murray et al., 2011; Lind et al., 2017). Selection pressures that may have an influence on the composition and diversity of BGCs in introduced populations are differences in ecological interactions, such as competition with novel microbial communities or changes in resource availability. Additionally, introductions of *S. luteus* to new environments often involved small founder populations with limited genetic diversity (Pildain et al., 2021; Ke et al., 2024, unpublished). Genetic drift and founder effects in these introduced populations could result in the loss or reduction of gene groups within BGCs over time, leading to lower gene group counts compared to native populations.

One limitation of this study is that genome coverage varied widely, and uniformly higher coverage would have likely produced higher quality assemblies and more robust BGC predictions. We also currently lack a chromosome-level reference genome assembly for *Suillus luteus*, which would also make genome assembly and gene predictions more reliable. As for sampling, locations were unevenly represented, with some populations much more highly represented than others, within both the native and introduced ranges. Additionally, the fungal version of AntiSMASH used here (fungiSMASH) was developed primarily with bacteria and Ascomycete non-mycorrhizal fungi (Almeida et al., 2022; Navarro-Munoz and Collemare, 2022), and shows a bias toward known BGC classes with predicted core enzymes, potentially limiting the scope of our results.

Our results provide new insights into how EcM fungus BGCs are evolving among native and introduced populations, revealing that BGC diversity varies on the gene scale while displaying conserved functional diversity overall. Introduced populations originating from the Central Europe clade show reduced diversity, while populations from Asia and Northern Europe exhibit unique BGC compositions, potentially indicating a cryptic species. Clan grouping revealed shared BGC gene groups and genetic diversification through changes in gene size, copy number, group diversity, and sequence similarity. Further functional annotations of BGCs within clans and lab validation of gene clusters and their chemicals are needed to confirm our findings and their potential applications. Given their structural diversity and widespread presence, analyzing natural products and their underlying BGCs is essential for understanding microbial interactions, involved organisms, and their environmental impacts.

## Supporting information

Supplementary Table 1. Genome assembly stats

Supplementary Table 2. Metadata, BGC predictions, GCFs, and clans for 258 genome

## 6. Acknowledgements

NSF grant #1953299

NSF grant # 1953405

## 7. Supplementary Material

**Supplementary Figure 1.**
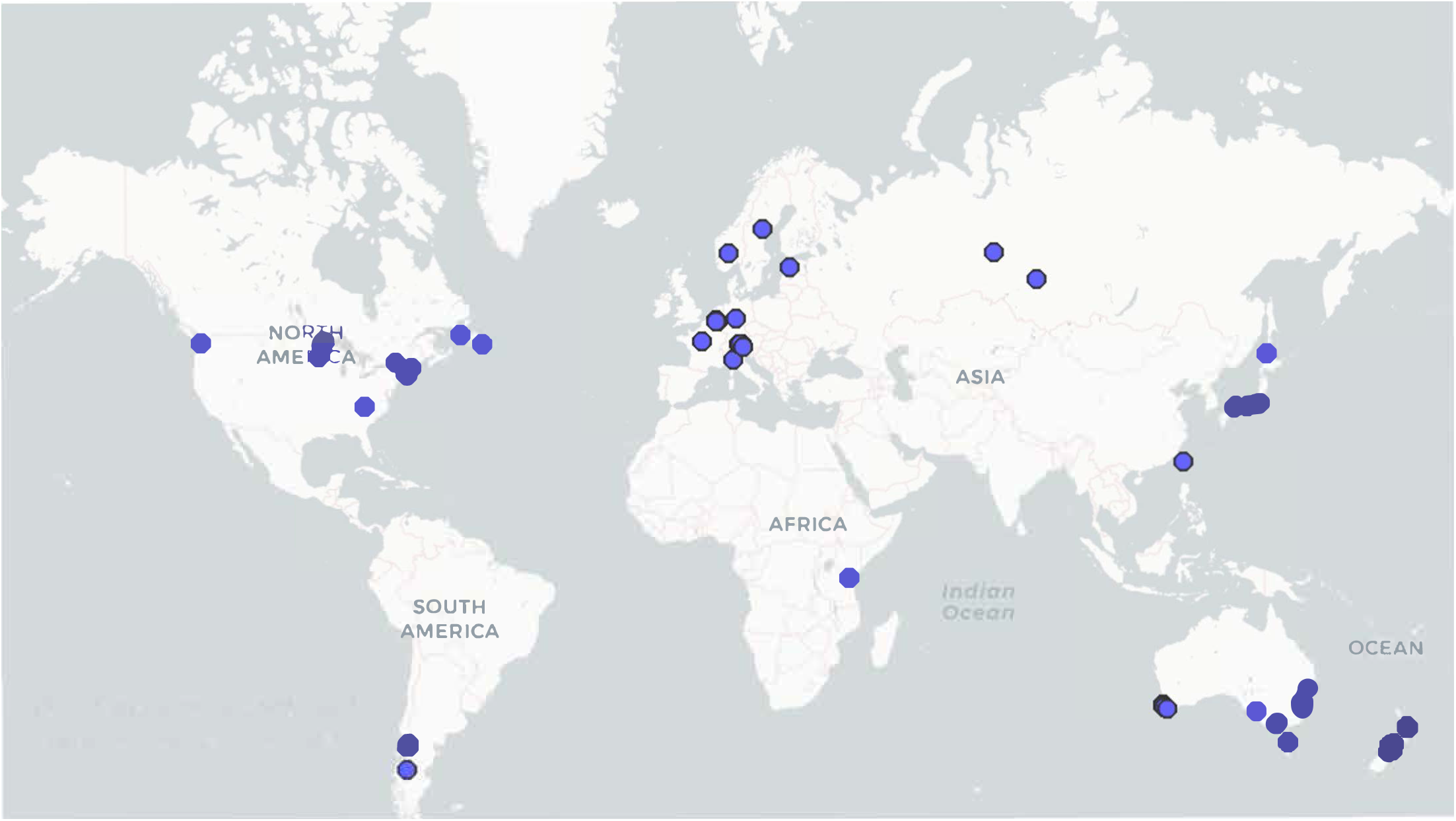
Map of genome sample locations

Supplementary Table 1. Genome assembly stats

Supplementary Table 2. Metadata, BGC predictions, GCFs, and clans for 258 genomes

